# Anatomo-functional correlates of auditory development in infancy

**DOI:** 10.1101/585448

**Authors:** Parvaneh Adibpour, Jessica Lebenberg, Claire Kabdebon, Ghislaine Dehaene-Lambertz, Jessica Dubois

## Abstract

Brain development incorporates several intermingled mechanisms throughout infancy leading to intense and asynchronous maturation across cerebral networks and functional modalities. Combining electroencephalography (EEG) and diffusion magnetic resonance imaging (MRI), previous studies in the visual modality showed that the functional maturation of the event-related potentials (ERP) during the first postnatal semester relates to structural changes in the corresponding white matter pathways. Here we aimed to investigate similar issues in the auditory modality. We measured ERPs to syllables in 1-to 6-month-old infants and analyzed them in relation with the maturational properties of underlying neural substrates measured with diffusion tensor imaging (DTI). We first observed a decrease in the latency of the auditory P2, and a decrease of diffusivities in the auditory tracts and perisylvian regions with age. Secondly, we highlighted some of the early functional and structural substrates of lateralization. Contralateral responses to monoaural syllables were stronger and faster than ipsilateral responses, particularly in the left hemisphere. Besides, the acoustic radiations, arcuate fasciculus, middle temporal and angular gyri showed DTI asymmetries with a more complex and advanced microstructure in the left hemisphere, whereas the reverse was observed for the inferior frontal and superior temporal gyri. Finally, after accounting for the age-related variance, we correlated the inter-individual variability in P2 responses and in the microstructural properties of callosal fibers and inferior frontal regions. This study combining dedicated EEG and MRI approaches in infants highlights the complex relation between the functional responses to auditory stimuli and the maturational properties of the corresponding neural network.

## Introduction

The development of the human brain is characterized by a period of fast changes during the first two years after birth followed by a slower but persistent progression until early adulthood. This protracted maturation enables children to acquire sensorimotor capacities and cognitive skills in connection with their environment. The multiple neural changes occurring during childhood are still incompletely understood, but two major phenomena supporting behavioral changes have been highlighted since the beginning of the 20th century by *post-mortem* examinations (Flechsig, 1920; Yakovlev. & Lecours., 1967): synaptogenesis-pruning cycles in the gray matter, mostly in the cortex, and the myelination of white matter fibers. Although these studies remained limited by the small sample size, brain anomalies and the impossible matching with infants’ functional development, they underscored the heterogeneity of the maturational patterns, with different timelines across regions, networks and functional systems.

With the rise of neuroimaging, it is now possible to examine brain development in healthy infants and over larger groups. Particularly structural Magnetic Resonance Imaging (MRI) studies have started to extend the former *post-mortem* studies, confirming the lengthy and heterogeneous changes occurring in the human brain in terms of cortical thickness (Li, Lin, Gilmore, & Shen, 2015; Sowell et al., 2003), surface (Yakovlev & Lecours, 1967) and microstructure (Leroy et al., 2011), as well as white matter maturation (Dubois, Dehaene-Lambertz, et al., 2014). In the cortical gray matter, indirect markers of the growth of dendritic arborization and synaptogenesis (Huttenlocher and Dabholkar, 1997) and of the myelination of intra-cortical fibers (Thomas et al., 2000) can be measured through relaxometry MRI (Leroy et al., 2011), diffusion MRI (Ball et al., 2013; Batalle et al., 2019) or both (Lebenberg, Mangin, et al., 2019). In particular, an asynchrony in microstructural maturation between primary and nonprimary auditory regions has been observed during the preterm period, based on parameters of diffusion tensor imaging (DTI) (Monson et al., 2018). In the white matter, the progressive axons ensheathment by myelin, the increasing lipid content and the fiber compactness constrain the diffusion of water molecules, leading to decreases in DTI diffusivities and an increase in fractional anisotropy in most tracts throughout infancy (Dubois, Hertz-Pannier, Dehaene-Lambertz, Cointepas, & Le Bihan, 2006; Huppi et al., 1998).

These structural changes have functional consequences that can be assessed in infants with electro-or magneto-encephalography (EEG, MEG) (Dubois, Adibpour, Poupon, Hertz-Pannier, & Dehaene-Lambertz, 2016). The topography and latencies of event-related potentials (ERPs) are changing, especially during the first postnatal year. For instance, in the visual system, the latency of the visual P1 (the first positive component following the presentation of a visual stimulus) decreases from ~300 ms at birth to ~100-120 ms (close to the adults’ values) about 3 months later (McCulloch, Orbach, & Skarf, 1999). Since myelination accelerates the conduction of neural transmissions, we expected that DTI parameters reflecting the maturation of white matter fibers would correlate with the speed derived from the latencies of ERP components. We confirmed this hypothesis in two successive studies in infants showing that the speed of the visual P1 was correlated with the transverse diffusivity measured in the optic radiations, and the speed of inter-hemispheric visual transfer with that of the splenial callosal fibers (Adibpour, Dubois, & Dehaene-Lambertz, 2018; Dubois et al., 2008).

Here, our goal was to investigate whether these results can be generalized to another sensory modality-the auditory system, and to further add another measure of maturation: the microstructural changes in the cortical gray matter where neural computations are performed, and which might be relevant to explain some variability across infants. The auditory system has a different developmental timeline compared with the visual system. Because of the noisy womb environment, the auditory cortex is already stimulated before birth, whereas visual stimulations are largely different *in-* and *ex-utero.* From the seminal work of Yakovlev and Lecours (1967) on the myelination of white matter tracts, it appears that acoustic radiations start to myelinate shortly after birth and continue slowly throughout the first 3 years, whereas this process occurs much more rapidly over the first postnatal semester for optic radiations. Consisting of a broad positive peak (P2) followed by a broad negative peak (N2), the infant’s auditory ERPs show slower changes than the visual ERPs: from birth to one year of age, latencies decrease from ~300 ms to ~150 ms for the P2, and from ~530 ms to ~300ms for the N2 (Barnet, Ohlrich, Weiss, & Shanks, 1975; Kushnerenko et al., 2002; Novak, Kurtzberg, Kreuzer, & Vaughan, 1989; Wunderlich & Cone-Wesson, 2006). Protracted changes are also observed throughout childhood (Shafer, Yan, & Wagner, 2015), notably with the appearance of earlier peaks (P1, N1) which are barely identifiable before 4-5 years of age (Lippé, Kovacevic, & McIntosh, 2009; Ponton, Eggermont, Kwong, & Don, 2000).

With regard to structure-function relationships, only one study has investigated this issue in the auditory domain from childhood to adolescence, relating DTI parameters in acoustic radiations and the latency of M100 auditory response evaluated with MEG (Roberts et al., 2009). However, the observed relationship did not survive when the children’s age was first entered in the statistical model, contrarily to our reports for the infant’s visual system (Adibpour, Dubois, & Dehaene-Lambertz, 2018; Dubois et al., 2008). This result suggests that the links between the structural and functional properties as we can measure them, might be not as straightforward for the auditory modality as for the visual one. This issue thus needs to be further investigated.

A second important characteristic of the human adult auditory system is that it is lateralized, some acoustic features being better processed by the left hemisphere, some by the right (Boemio, Fromm, Braun, & Poeppel, 2005). Signs of functional lateralization are seen early on. Preterms at 6 month of gestation and listening to speech syllables display faster and more sustained BOLD responses over the left posterior temporal region than the right, escaping the general pattern of larger right responses measured over the other regions (Mahmoudzadeh et al., 2013). Dehaene-Lambertz et al (2010) reported larger activations in the left than right *planum temporale* for speech, whereas activations were symmetric for piano music in 2-3-month-old infants (Dehaene-Lambertz et al., 2010). By contrast, Perani et al (2011) reported larger activations in a right than left sphere defined around Heschl’s gyrus in neonates listening to speech, but the whole-brain analysis showed mainly bilateral activations and even left activations when hummed and flattened speech were compared (Perani et al., 2011). Finally, Shultz et al (2014) observed an increase with age in the difference of activations for speech and biological non-speech sounds in the left temporal lobes, when all responding voxels were pooled (Shultz, Vouloumanos, Bennett, & Pelphrey, 2014). This suggests an increase in the left hemispheric bias for speech perception during infancy. Besides, we reported early hemispheric asymmetries of the language network at the structural level, in terms of sulcation (Dubois et al., 2010; Glasel et al., 2011), cortical maturation (Leroy et al., 2011; Rolland et al., 2019) and white matter architecture (Dubois et al., 2009; Dubois, Poupon, et al., 2016). Therefore, here we aimed to evaluate the relationships between early functional and anatomical markers of lateralization.

In this study, we used complementary EEG and DTI measures to study the maturation of the auditory network in infants during the first postnatal semester, when changes are intense and when structure-function relationships have already been demonstrated in the visual domain (Adibpour, Dubois, & Dehaene-Lambertz, 2018; Dubois et al., 2008). After analyzing how each of the functional and structural indices evolves with infants’ age, we analyzed hemispheric differences and investigated the extent to which these complementary measures of maturation are inter-related in the auditory domain.

## Material and Methods

### 1. Subjects

EEG: We tested a first group of 23 infants in a binaural auditory paradigm (postnatal age between 5 and 21.4 weeks, 10 girls and 13 boys), and a second group of 19 infants in a monaural auditory paradigm (age between 6.7 to 28.7 weeks, 6 girls and 13 boys).

MRI: Diffusion MRI data were acquired in a group of 22 infants (age between 5.9 to 22.4 weeks, 9 girls and 13 boys). Analyses over cortical regions only included 21 of them because anatomical images required for segmentation could not be acquired for one infant.

Some infants participated to both EEG (monaural paradigm) and MRI exams, a few days apart: n=16 with EEG and diffusion MRI data (5 girls and 11 boys); n=15 with EEG, diffusion and anatomical MRI data (4 girls and 11 boys).

All infants were considered as typical subjects (born full-term, with no reported history of medical problems). The study was approved by the regional ethical committee for biomedical research. All parents were informed about the content and goals of the experiments, and gave written informed consent.

### 2. EEG study

#### 2.1 Experimental paradigms

##### Stimuli

Two consonant–vowel syllables (/pa/ and /ta/) with neutral intonation were produced by two female speakers and matched for intensity and total duration (200 ms).

##### Binaural Paradigm

In each trial, syllables were presented by series of four, with an interval of 600 ms between the stimuli onsets and 2.2 s of silence following the fourth syllable onset and before the onset of the next trial (leading to a total duration of each trial of 4 s). At each trial, the syllable (phoneme /pa/ or /ta/, speaker) was randomly chosen among the four possibilities, and presented three times. The last syllable determined the type of trials: it was either similar to the first three (standard trials), or there was a change of phoneme (deviant phoneme trials) or a change of voice (deviant voice trials). A maximum of 192 trials was presented in 4 blocks of 48 trials (corresponding to 3 types of trials x 2 phonemes x 2 voices x 4 repetitions).

##### Monaural Paradigm

Small earphones were placed on the infant’s ears and maintained in place by the EEG net. The paradigm was similar to the binaural paradigm except that the first three syllables were presented in a single ear, and the last syllable was either played in the same or in the other ear. The initial ear side was kept constant during a block. 192 trials were presented in 4 blocks (2 blocks per ear side) of 48 trials (corresponding to 2 ear sides for the last syllable x 3 types of trials x 2 phonemes x 2 voices x 2 repetitions).

#### 2.2. EEG data acquisition

An EEG net comprising 128 electrodes (EGI, Eugene, USA) with a reference located on the vertex was placed on infant’s head relative to anatomical markers on the scalp. EEG was continuously digitized at a sampling rate of 250 Hz during the whole experiment (net amp 200 system EGI, Eugene, USA). The infants were sitting on the parent’s laps in a shielded room. They may have fallen asleep or remained awake during the experiment. In the latter case, one experimenter was showing them interesting objects and cartoon images to keep them quiet. The experiment was stopped before the end if infants became restless.

#### 2.3. EEG pre-processing

EEG recordings were band-pass filtered between 0.5 and 20 Hz using a zero-phase lag filter, and were further processed using MATLAB toolboxes: EEGLAB (Delorme & Makeig 2004) and Brainstorm (Tadel et al. 2011). For both paradigms, the signal was segmented into epochs of 3700 ms [-200, +3500] ms relative to the onset of the first syllable at each trial. Channels contaminated by movement or eye artifacts were automatically rejected on a trial by trial basis, based on amplitude variations inside each epoch: for each channel, an epoch was rejected when the fast average amplitude exceeded 250 μv, or when deviation between fast and slow running averages exceeded 150 μv. Channels were rejected if they were marked as bad in more than 70% of the trials, and trials were rejected if more than 50% of electrodes were marked as bad. Recordings were then re-referenced by subtracting the average activity of all channels over the brain to obtain average-reference recordings, then baseline-corrected by [-200 0] ms time-window before the onset of the first syllable.

We computed auditory-evoked responses by averaging all correct trials in each condition. For the binaural experiment, we kept on average a total of 140 out of 192 trials. For the monaural experiment, we averaged all correct trials for each ear side independently and kept on average a total of 68/69 out of 96 trials for the left/right ear respectively. Here, we focused on brain responses to the first syllable only, since it showed the highest signal-to-noise ratio and so the most reliable identification of the ERP components.

#### 2.4. Identification of auditory evoked potentials

To measure the latency of auditory evoked responses, we focused on the positive pole of the auditory response which is extended more laterally compared to the more medial negative pole (see Figure 1). We identified two symmetrical clusters of electrodes over the left and right sides on the topography of the grand average performed across each group. For the binaural paradigm, we considered 27 electrodes over each hemisphere, covering the fronto-temporal regions and extending over T1-F7 on the left side (T2-F8 on the right) (Figure 1. a). For the monaural experiment, we identified two symmetrical clusters of 15 electrodes, around T7/T8 extending anteriorly up to F3/F4, and posteriorly down to P7/P8 (Figures 1. b). For each infant, we averaged the activity over each cluster of electrodes.

**Figure 1:**
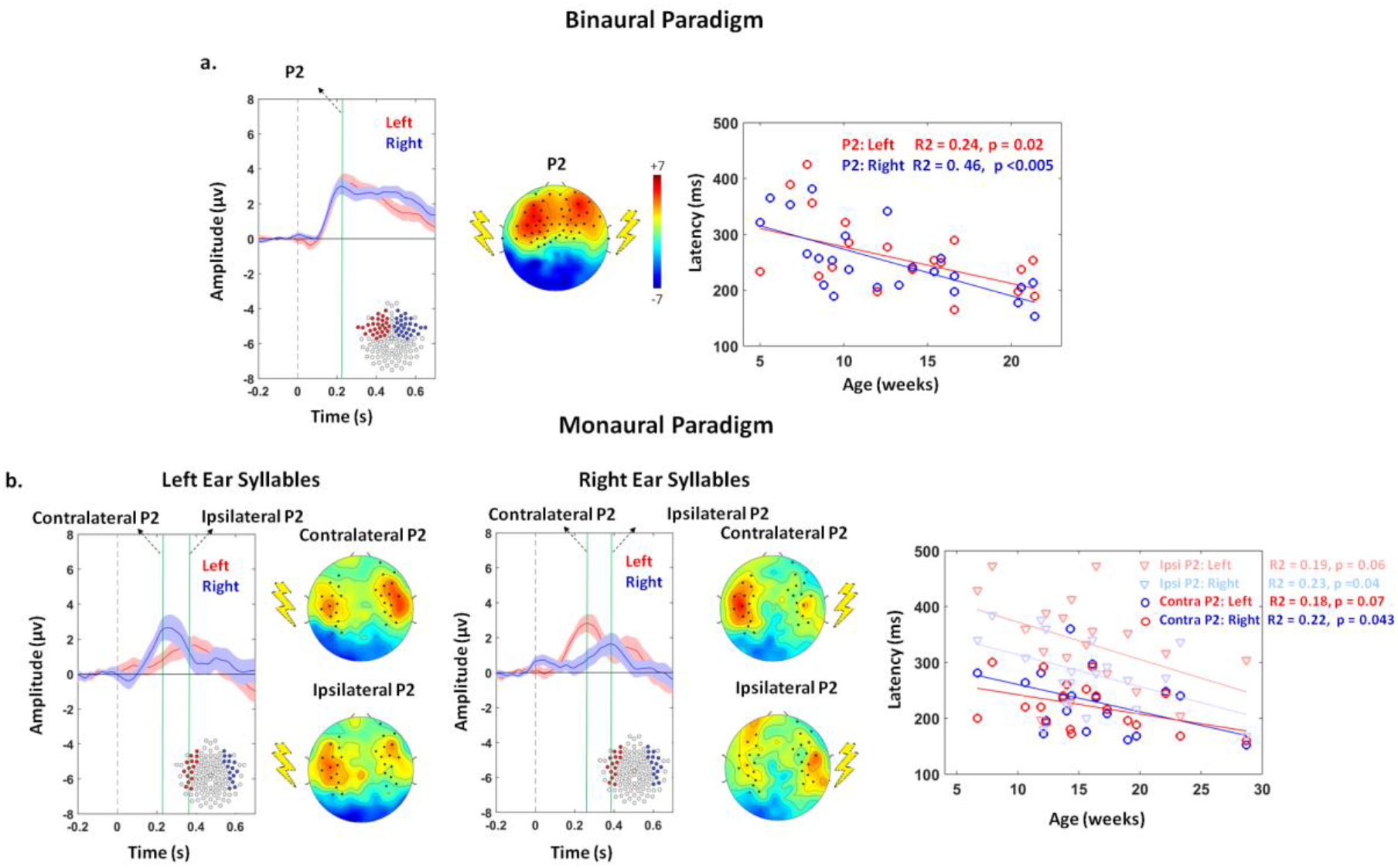
Time course of auditory evoked responses and age-related decrease in P2 latency. Left panels: Grand average ERPs over the groups in response to the first syllable presented either binaurally (a: n=23), or monaurally in the left and right ears (b: n=19). The grand-averages computed over the highlighted clusters of electrodes are displayed (red=left side, blue=right side). The voltage topography at P2 peak is presented next to the time course plot. For monaural syllables, topographies are presented at the time of the contralateral and ipsilateral P2 peaks. Time zero marks the syllable onset. Right panels: In each infant, the peak of P2 response was identified over the left (red) and right (blue) clusters for syllables presented binaurally, or monaurally. In b. darker colors correspond to the contralateral responses, and light colors to the ipsilateral ones. The latency of each P2 response decreases with age.

In this study we focused on P2, which is the most prominent auditory response in the first semester after birth (Wunderlich & Cone-Wesson., 2006) and can be more easily identified in each infant. We identified P2 component as the first distinguishable positive peak, appearing between 150-450 ms in the time course of auditory evoked potentials. In each infant, we measured P2 latency at the peak, and computed its amplitude over a 50 ms time window centered on the peak.

### 3. MRI study

#### 3.1. MRI acquisition and pre-processing

Acquisitions were performed during spontaneous sleep in a 3T MRI scanner (Tim Trio, Siemens Healthcare, Erlangen, Germany), equipped with a whole-body gradient (40mT/m, 200T/m/s) and a 32-channel head coil. To minimize specific absorption rate (SAR) and noise exposure, we used radio-frequency (RF) impulsions with “no SAR”, and “whisper” gradient mode when possible. Infants’ ears were protected from the noise using headphones placed over their ears during the acquisition.

Anatomical T2-weighted (T2w) images were also acquired in infants using a 2D turbo spin echo sequence (spatial resolution = 1}1}1.1mm3), providing the best grey / white matter contrast at these ages (Dubois et al. 2014). For DTI and tractography purposes, a diffusion-weighted (DW) spin-echo single-shot EPI sequence was used, with parallel imaging (GRAPPA reduction factor 2), partial Fourier sampling (factor 6/8) and monopolar gradients to minimize mechanical and acoustic vibrations. After the acquisition of the b=0 volume, diffusion gradients were applied along 30 orientations with b=700s.mm-2. Fifty interleaved axial slices covering the whole brain were acquired with a 1.8mm isotropic spatial resolution (field of view = 230}230 mm^2^, matrix = 128}128, slice thickness = 1.8mm, TE = 72ms, TR = 10s), leading to a total acquisition time of 5min40s which was reasonably short for unsedated infants. Motion artifacts were corrected on DW images using PTK/Connectomist software (Dubois, Kulikova, et al., 2014; Duclap et al., 2012).

#### 3.2. Identification of white matter tracts and microstructure characterization

Probabilistic tractography was based on a 2-crossing-fiber diffusion model using FSL software (Behrens et al. 2007) over individual brain masks. Using individual seed regions, the following fiber tracts were identified from the auditory network: 1. left and right acoustic radiations (AR) (by locating seeds at the level of medial geniculate nucleus in thalamus, and temporal regions around Heschl’s gyrus); 2. auditory fibers of the corpus callosum (ACC) (by locating the seeds in the left and right auditory regions) 3. left and right arcuate fascicles (AF) (by locating seeds at the superior parietal part of the tract, and at parieto-temporal junction at the level of the arcuate loop).

Following the estimation of the diffusion tensor, DTI parameters were quantified in each voxel of each infant brain. For each tract, averaged parameter X was calculated over the tract voxels by taking into account fiber density on the tract density map (Hua et al. 2008):

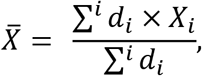

where i denotes the tract voxels, d_i_ is the fiber density at voxel i, and X_i_ is the value of the DTI parameter at voxel i. In this study, we focused on transverse diffusivity (λ_⊥_) as it was shown to be the best DTI marker of white matter myelination according to previous studies in animal models of demyelination (Song et al., 2003; Song et al., 2005) and infant studies of white matter maturation (Dubois, Dehaene-Lambertz, et al., 2014). Moreover, we previously showed that a higher speed of visual responses is related to a lower transverse diffusivity in the corresponding tracts (Adibpour, Dubois, & Dehaene-Lambertz, 2018; Dubois et al., 2008).

#### 3.3. Identification of cortical regions and microstructure characterization

In each infant, anatomical images were used to extract the cortical surface through a semi-automatic procedure (Dubois et al., 2019). An individual atlas of cortical parcels defined on a 10-weeks-old infant (Kabdebon et al., 2014) was projected on individual surfaces using a reliable inter-individual registration based on cortical sulci and ribbons (Lebenberg, Labit, et al., 2019). Here we focused on the auditory-related perisylvian regions: Heschl’s gyrus (HG), *Planum Temporale* (PT), superior and medial temporal gyrus (STG, MTG), inferior frontal gyrus (IFG), and angular gyrus (AG).

We mapped DTI parameters over the cortical ribbon in each infant using a precise intraindividual registration correcting for geometric distortions (Lebenberg, Mangin, et al., 2019; Lebenberg et al., 2015). To specifically identify cortical measures, we took advantage of the fact that longitudinal diffusivity (λ_//_) displays minimal values in the cortex compared with surrounding tissues regardless of the maturational stage (Lebenberg, Mangin, et al., 2019). We computed median DTI parameters over each cortical region. We focused on λ_//_ to characterize the cortical microstructure because it reliably decreases with increasing maturation over this developmental period (Lebenberg, Mangin, et al., 2019; Rolland et al., 2019).

### 4. Statistical analyses

#### 4.1. Dependences related to age, hemisphere, side and region

We first evaluated the factors that might modulate the ERPs functional characteristics. We performed analyses of covariance (ANCOVA) on the P2 amplitudes and latencies for the binaural (n=23) and monaural (n=19) paradigms independently, with age as between-subjects covariate and hemisphere (left/right) as a within-subject factor. For the monaural paradigm, we entered a supplementary within-subject factor for the response side (contralateral/ipsilateral side relatively to the ear of syllable presentation).

Second, we assessed similar dependences for DTI structural parameters (transverse diffusivity for each white matter tract: n=22, longitudinal diffusivity for each cortical region: n=21). With the diffusivity as dependent variable, we performed ANCOVA with age as between-subjects covariate, tract/region and hemisphere (left/right) as within-subject factors.

In all these analyses, hemispheric asymmetries were tested through the factor “hemisphere”. When the tract/region x hemisphere interaction was significant, post hoc analyses were performed using paired t-tests. To limit the number of comparisons, we focused on the most relevant pairs (i.e. across different tracts/regions within each hemisphere, and across homologous tracts/regions of the two hemispheres). For the post hoc analyses, p-values were corrected for the number of comparisons using FDR approach.

#### 4.2. Relationships between the functional markers of maturation

We tested whether asymmetry indices (defined as (L-R)/(L+R) with left L and right R values for amplitude or latency) were related between contralateral and ipsilateral responses using Pearson correlations (these measures did not depend on the infants’ age).

#### 4.3. Relationships between the structural markers of maturation

As we considered auditory white matter tracts and perisylvian cortical regions that all belong to the same functional network, we further investigated whether these different neural substrates have related DTI microstructural changes over this developmental period, while controlling for common dependencies on infants’ age. As previously, we evaluated relationships for the most relevant comparisons, i.e. across i) pairs of tracts/regions in each hemisphere (n=22/21), ii) homologous tracts/regions of left and right hemispheres (n=22/21), and iii) pairs of tracts and regions in each hemisphere (n=21). For each paired comparison, we used partial correlations over the group, controlling for age. P-values were corrected for the number of comparisons in each hemisphere condition (e.g. all pairs of tracts in the left hemisphere) using FDR approach.

We also tested whether asymmetry indices in diffusivity parameters were related to each other across tracts/regions using Pearson correlations (these measures did not depend on the infants’ age). Although no asymmetry index can be computed for callosal fibers, we included this tract as it is since it might interplay with other asymmetries for the development of lateralization.

#### 4.4. Relationships between the structural and functional markers of maturation

Finally, we aimed to examine the relationships between functional and structural markers of maturation in infants who had both EEG (monaural paradigm) and DTI datasets. In the following analyses, we considered left and right measures independently when hemispheric asymmetry was observed based on previous analyses. Otherwise, the averaged value was used in order to limit the number of comparisons.

Following our studies of the visual modality (Dubois et al. 2008, Adibpour, Dubois & Dehaene-Lambertz, 2018), we first compared the maturational properties of auditory-evoked responses and of white matter tracts (n=16). As latencies are assumed to depend both on the fibers myelination and on the distance the neural signal travels, we considered a marker of speed, defined as the ratio between the fiber length and latency. We approximated the length of auditory pathways by the distance between the two ears measured on MRI images for each infant. We then analyzed whether the speed of P2 responses (averaged contralateral, left and right ipsilateral) was related to the transverse diffusivity in tracts that might contribute to neural propagation in this network (left and right acoustic radiations, arcuate fascicles, and auditory callosal fibers). Note that contrarily to our visual study (Adibpour, Dubois & Dehaene-Lambertz, 2018), we could not directly measure the inter-hemispheric transfer of responses in the auditory modality, because responses to monaural stimuli originate from both contralateral and ipsilateral pathways and are observed quite simultaneously in both hemispheres (Adibpour, Dubois, Moutard, & Dehaene-Lambertz, 2018).

Since the microstructure of cortical regions might also contribute to the properties of P2 responses during infancy, we performed similar analyses considering longitudinal diffusivity in the different perisylvian regions (n=15). Following a recent study comparing the visual and auditory modalities in the aging population (Price et al., 2017), we primarily considered relationships with P2 latencies (results for P2 speeds are reported in supplementary information).

As previously, we performed partial correlations over the group, controlling for the infants’ age, and we corrected for multiple comparisons in each tested condition using FDR approach.

Furthermore, we investigated whether the asymmetry indices in functional and structural markers were related using Pearson correlations, and considering i) P2 speeds vs white matter tracts (n=16) and ii) P2 latencies vs cortical regions (n=15).

## Results

### Development and asymmetry of P2 functional responses

Figure 1 presents the ERP topographies in each experimental task. The positive pole of the P2 component extended more over the frontal midline areas in the case of binaural compared with monaural stimulation. While we observed brain responses on both brain sides following monaural stimulation, the contralateral responses were larger and more extended than the ipsilateral ones, whatever the side of stimulation.

#### P2 amplitude

ANCOVA (Table 1a) revealed a marginally significant effect of hemisphere due to a larger amplitude in the left than right hemisphere when stimulation was binaural, but no effect of age nor interaction between age and hemisphere. Regarding monaural stimulation (Table 1b), only a marginally significant effect of response side was observed due to larger amplitude in the contralateral than ipsilateral side. Asymmetry indices for contralateral and ipsilateral amplitudes (Figure 2a) were not related to each other (r=0.06, p=0.786).

**Figure 2:**
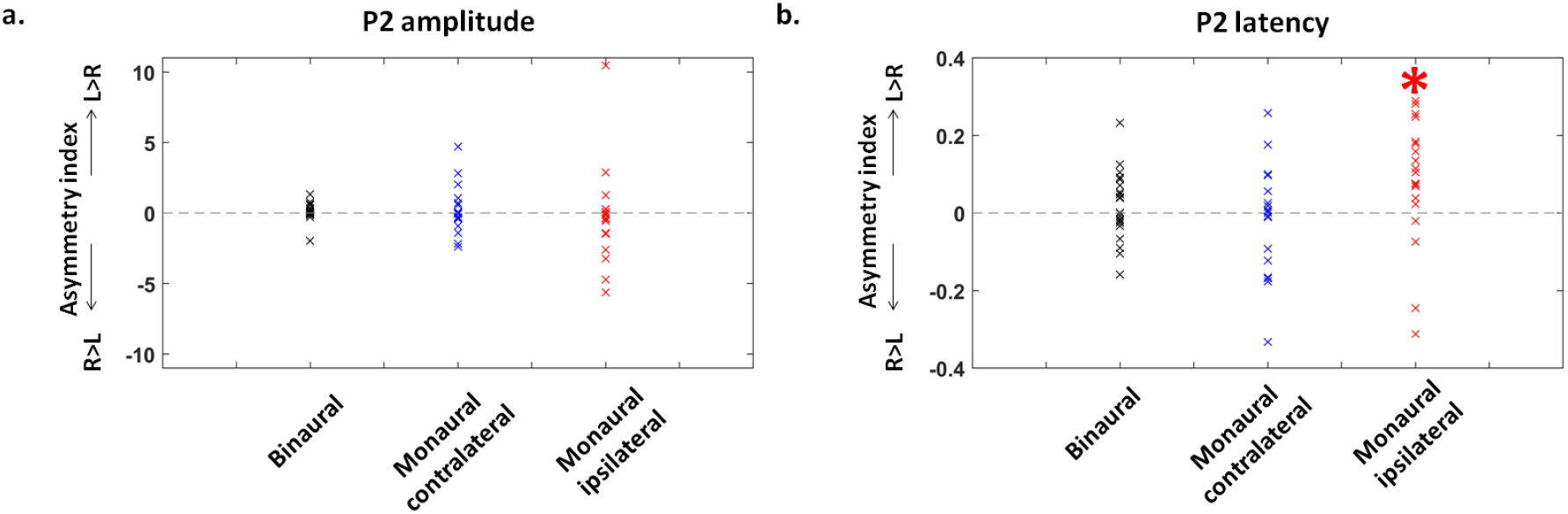
Hemispheric differences in the amplitude and latency of P2 peaks. Asymmetry indices (Left-Right)/(Left+Right) for P2 amplitude (a) and latency (b) in the different paradigms: binaural (n=23) and monaural syllables (n=19: contralateral and ipsilateral responses). The latency of ipsilateral responses to monaural syllables was significantly slower over the left than right cluster, in other words the ipsilateral response for a left ear syllable was slower than for a right ear syllable (*: p <0.05). Statistical analyses are detailed in Table 1.

**Table 1:** Summary of ANCOVA results for the amplitude and latency of P2 responses to binaural (a) monaural (b) syllables. Amplitudes and latencies were considered as the dependent variable in separate ANCOVA models with age, hemisphere (left/right) and response side (contralateral/ipsilateral in the case of monaural syllables) entered as independent variables (1 covariate and 2 within-subject factors). For responses to monaural syllables, further post hoc analyses were performed to compare pairs of P2 latencies using paired t-test comparisons (p-values were corrected for the number of comparisons using FDR approach: c=6). Significant (p<0.05) and marginally significant (p<0.1) comparisons are shaded in green and yellow.

| a. Binaural Presentation |  |
| --- | --- |
| Amplitude | Latency |
| age : $F(1,21) < 1$, $p = 0.494$ | age : $F(1,21) = 13.1$, $p = 0.002$ |
| hemisphere : $F(1,21) = 3.0$, $p = 0.095$ | hemisphere : $F(1,21) < 1$, $p = 0.385$ |
| age x hemisphere : $F(1,21) < 1$, $p = 0.380$ | age x hemisphere : $F(1,21) < 1$, $p = 0.385$ |
| b. Monaural Presentation |  |
| Amplitude | Latency |
| age : $F(1,17) = 1.6$, $p = 0.220$ | age : $F(1,17) = 15.2$, $p = 0.001$ |
| hemisphere : $F(1,17) < 1$, $p = 0.630$ | hemisphere : $F(1,17) = 1.7$, $p = 0.210$ |
| response side : $F(1,17) = 3.4$, $p = 0.083$ | response side : $F(1,17) = 44.5$, $p < 0.001$ |
| age x hemisphere : $F(1,17) < 1$, $p = 0.377$ | age x hemisphere : $F(1,17) < 1$, $p = 0.943$ |
| age x response side : $F(1,17) < 1$, $p = 0.923$ | age x response side : $F(1,17) < 1$, $p = 0.393$ |
| hemisphere x response side : $F(1,17) < 1$, $p = 0.815$ | hemisphere x response side : $F(1,17) = 10.7$, $p = 0.004$ |
| age x hemisphere x response side : $F(1,17) < 1$, $p = 0.926$ | age x hemisphere x response side : $F(1,17) < 1$, $p = 0.530$ |
|  | Posthoc comparisons |
| | contra left vs contra right : $t(1,18) < 1$, $p = 0.495$ |
| | contra left vs ipsi left : $t(1,18) = -6.3$, $p < 0.001$ |
| | contra left vs ipsi right : $t(1,18) = -3.4$, $p = 0.004$ |
| | contra right vs ipsi left : $t(1,18) = -4.6$, $p < 0.001$ |
| | contra right vs ipsi right : $t(1,18) = -4.0$, $p = 0.002$ |
| | ipsi left vs ipsi right : $t(1,18) = 2.5$, $p = 0.028$ |

#### P2 latency

ANCOVA (Table 1a) showed a main effect of age (latency decreased with age: Figure 1a), but no difference between hemispheres, nor interaction between age and hemisphere when stimulation was binaural. For monaural stimulation (Table 1b), we also observed a main effect of age (latency decreased with age: Figure 1b), as well as a main effect of response side, with an interaction between side and hemisphere. Post hoc analyses (Table 1b) revealed that i) P2 latency was longer on the ipsilateral than contralateral side whatever the hemisphere, and ii) ipsilateral responses had longer latency in the left than right hemisphere, whereas no hemispheric difference was seen for contralateral responses. Asymmetry indices in contralateral and ipsilateral latencies (Figure 2b) were not related to each other (r=0.19, p=0.445).

### Development and asymmetry of auditory structural substrates

#### White matter

Tracts were precisely reconstructed in each infant with tractography (Figure 3a). A first ANCOVA on transverse diffusivity averaged over left and right acoustic radiations, arcuate and auditory fibers of the corpus callosum (Table 2a) revealed a main effect of age (diffusivity decreased with age: Figure 3a), as well as a main effect of tract but no interaction age x tract. A second ANCOVA separating the left and right AR and AF (Table 2a) confirmed these effects and showed a main effect of hemisphere, but no interaction of age with the other factors (2-way and 3-way interactions were not significant). In other words, the degree of tract asymmetries was not changing with age during this period. Then, we performed selected post hoc tests to compare the different tracts within each hemisphere, as well as left vs. right homologous tracts (Table 2a). Within a hemisphere, tracts demonstrated significant microstructural differences following a gradient of λ_⊥_ values (from higher to lower ones: AF > AR > ACC, Figure 3a). Across hemispheres, λ_⊥_ was higher in the right AR and AF relatively to their left homologs (Figure 4a). Partial correlations of λ_⊥_ values controlling for age dependences revealed that despite their microstructural differences, several tracts shared common maturational patterns (Sup Table 1a): homologous tracts, left AR with left AF, and bilateral AF with ACC. This suggested that most pathways are maturing in synchrony despite being at different maturational stages. Yet, the AR and AF asymmetry indices were not related to each other, nor with the ACC microstructure (in all the 3 comparisons: r<0.20, p>0.89 after correction for multiple comparisons).

**Figure 3:**
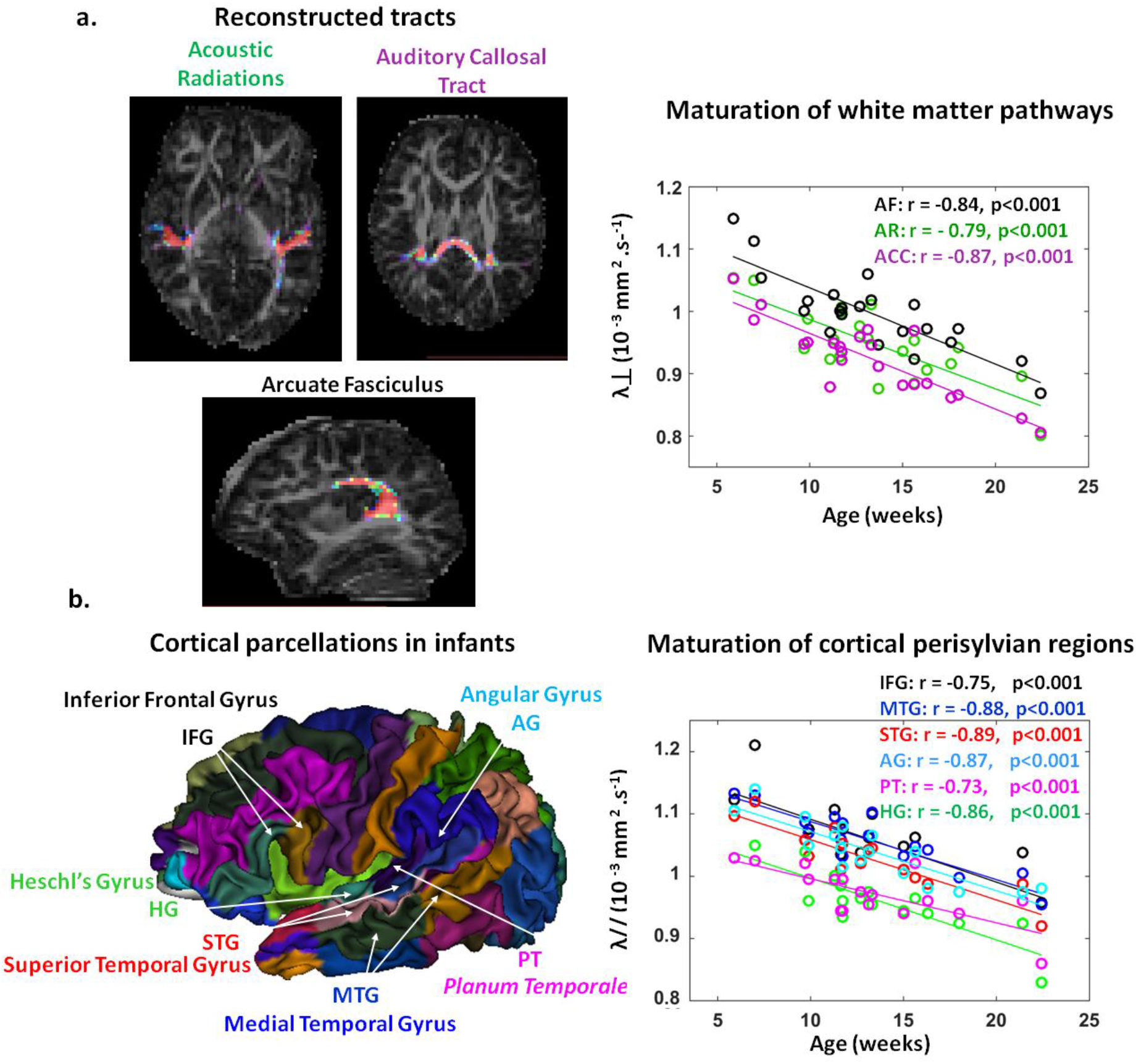
Maturation of the linguistic pathways and perisylvian regions quantified with diffusion MRI. a: Left panel: Example of tract reconstructions in a 12-week-old infant superposed on the map of DTI anisotropy: acoustic radiations (AR), auditory fibers of the corpus callosum (ACC) and arcuate fasciculus (AF). Right panel: Transverse diffusivity (λ_⊥_) plotted as a function of infants’ age in each tract (left and right values are averaged). Major age-related decreases are observed (n=22). b: Left panel: 3D parcellation atlas of a 10-week-old infant showing the studied regions: Heschl’s gyrus (HG), *planum temporale* (PT), superior and middle temporal gyri (STG, MTG), angular and inferior frontal gyri (AG, IFG). Right panel: Longitudinal diffusivity (λ_||_) plotted as a function of infants’ age in each region (left and right values are averaged). Major age-related decreases are observed (n=21).

**Table 2:** Summary of ANCOVA results for DTI parameters in white matter tracts (a) and perisylvian cortical regions (b). Transverse and parallel diffusivities were considered as the dependent variables in the ANCOVA models, with age, tract/region, hemisphere (left/right) entered as independent variables (1 covariate and 2 within-subject factors for the models of bilateral tracts and perisylvian regions). An additional ANCOVA is presented for combined left and right tracts (i.e. averaged transverse diffusivity over left and right acoustic radiations AR and arcuate fascicles AF) to include auditory fibers of the corpus callosum ACC without the hemisphere factor. We further selected post hoc analyses and performed comparisons between selected pairs of tracts/regions using paired t-tests: within each hemisphere (c=3 for tracts, c=15 for regions), and across hemispheres for homologous tracts/regions (c=2 for tracts, c=6 for regions) (p-values were corrected using FDR approach for the number of comparisons within each hemisphere condition). For the sake of simplicity, we present only the pairs of regions with the least significant results (all the other comparisons show more significance and can be derived from these pairs). Significant (p<0.05) and marginally significant (p<0.1) comparisons are shaded in green and yellow respectively. For the comparisons within each hemisphere, the pairs of tracts/regions were ordered to highlight the gradient of λ_⊥_/λ_//_ values (from higher to lower ones).

| Structural maturation |  |
| --- | --- |
| a. White matter tracts ($\lambda^+$) | b. Perisylvian regions ($\lambda_{II}$) |
| **ANCOVA with all tracts averaged over hemispheres** | age : F (1,19) = 51.5, p < 0.001 |
| age : F (1,20) = 62.6, p < 0.001 | region : F (5,95) = 101.5, p < 0.001 |
| tract : F (2,40) = 64.5, p < 0.001 | hemisphere : F (1,19) < 1, p = 0.717 |
| age x tract : F (2,40) < 1, p = 0.734 | age x region : F (5,95) = 1.1, p = 0.350 |
| **ANCOVA with only AR and AF** | age x hemisphere : F (1,19) < 1, p = 0.379 |
| age : F (1,20) = 47.6, p < 0.001 | region x hemisphere : F (5,95) = 7.5, p < 0.001 |
| tract : F (1,20) = 59.2, p < 0.001 | age x region x hemisphere : F (5,95) = 1.4, p = 0.233 |
| hemisphere : (1,20) = 14.8, p = 0.001 | **Posthoc comparisons** |
| age x tract : F (1,20) < 1, p = 0.457 | **Left Hemisphere** |
| age x hemisphere : F (1,20) < 1, p = 0.914 | IFG vs. MTG, t(1,20) = 2.3, p = 0.031 |
| tract x hemisphere : F (1,20) < 1, p = 0.673 | MTG vs. STG, t(1,20) = 3.6, p = 0.002 |
| age x tract x hemisphere : F (1,20) < 1, p = 0.796 | STG vs. AG, t(1,20) = 2.3, p = 0.033 |
| **Posthoc comparisons** | AG vs. PT, t(1,20) = 3.7, p = 0.002 |
| **Left Hemisphere** | PT vs. HG, t(1,20) = 3.2, p = 0.005 |
| AF vs. AR, t(1,21) = 7.9, p < 0.001 | **Right Hemisphere** |
| AF vs. ACC, t(1,21) = 9.4, p < 0.001 | MTG vs. IFG, t(1,20) = 2.2, p = 0.050 |
| AR vs. ACC, t(1,21) = 1.7, p = 0.096 | MTG vs. AG, t(1,20) = 1.5, p = 0.182 |
| **Right Hemisphere** | AG vs. IFG, t(1,20) = 1.2, p = 0.265 |
| AF vs. AR, t(1,21) = 5.6, p < 0.001 | IFG vs. STG, t(1,20) = 3.7, p = 0.002 |
| AF vs. ACC, t(1,21) = 15.9, p < 0.001 | AG vs. STG, t(1,20) = 6.0, p < 0.001 |
| AR vs. ACC, t(1,21) = 4.5, p < 0.001 | STG vs. PT, t(1,20) = 7.0, p < 0.001 |
| **Left vs Right tracts** | STG vs. HG, t(1,20) = 6.6, p < 0.001 |
| AF, t(1,21) = -2.1, p = 0.061 | PT vs. HG, t(1,20) < 1, p = 0.668 |
| AR, t(1,21) = -3.4, p = 0.004 | **Left vs Right regions** |
|  | IFG, t(1,20) = 3.9, p = 0.004 |
|  | MTG, t(1,20) = -2.1, p = 0.067 |
|  | STG, t(1,20) = 2.5, p = 0.044 |
|  | AG, t(1,20) = -4.6, p < 0.001 |
|  | PT, t(1,20) = 1.3, p = 0.252 |
|  | HG, t(1,20) < 1, p = 0.461 |

#### Grey matter

Perisylvian cortical regions were identified in each infant based on individual parcellation (Figure 3b). ANCOVA on longitudinal diffusivity (Table 2b) revealed a main effect of age (diffusivity decreased with age in all regions: Figure 3b), of region but no effect of hemisphere. The interaction region x hemisphere was significant, while the others were not. Selected post hoc analyses detailed the microstructural differences across regions, within each hemisphere and across hemispheres for homologous regions (Table 2b). They revealed a gradient of *λ_//_* values (from higher to lower ones: in the left hemisphere IFG > MTG > STG > AG > PT > HG; in the right hemisphere MTG ~/> AG ~ IFG > STG > PT ~ HG, Figure 3b). Regarding hemispheric asymmetries, higher diffusivity values were observed in the right AG and MTG relative to the left homologues, and in the reverse direction (L>R) for IFG and STG (Figure 4b). Despite these microstructural differences, partial correlations of *λ_//_* values controlling for age dependences revealed common maturational patterns across regions (Sup Table 1b). Except for PT and HG for which results were less clear-cut, homologous (left and right) regions were related, as well as each pair of regions (IFG, AG, STG and MTG) within each hemisphere. Besides the asymmetry indices across regions were not related to each other (in all the 15 comparisons: r–0.37, p>0.54 after correction for multiple comparisons).

**Figure 4:**
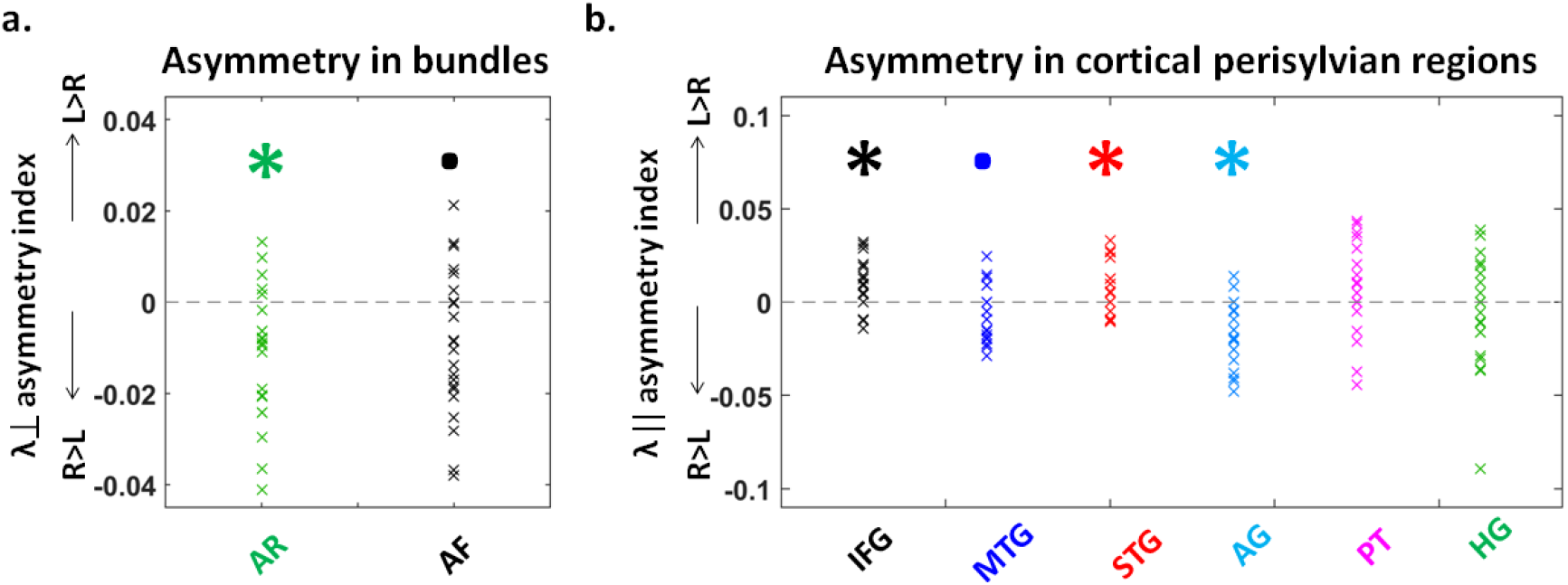
Hemispheric differences in the DTI properties of white matter tracts and cortical regions. Asymmetry indices (Left-Right)/(Left+Right) of DTI parameters in the white matter tracts (a, n=22) and in the cortical perisylvian regions (b, n=21). a. Lower transverse diffusivity (λ_⊥_) values in the left acoustic radiations (AR) and to a lesser extent in the left arcuate fasciculus (AF) compared to their right counterparts suggest a more mature microstructure in the left tracts. b. Similarly, the lower longitudinal diffusivity (λ_||_) in the left than right angular gyrus (AG) and middle temporal gyrus (MTG) to a lesser extent, suggests more mature regions in the left hemisphere. The reverse pattern was observed for the inferior frontal (IFG) and superior temporal (STG) gyri, suggesting a more mature microstructure in the right hemisphere for these regions (statistical analyses are detailed in Table 2; asterisks represent significant differences from zero at p<0.05, and dots a marginally significant trend at p<0.1).

We also evaluated the links between the different structural substrates composing the auditory network. Partial correlations between pairs of white matter tracts (considering λ_⊥_) and cortical regions (considering λ_//_)revealed reduced common maturational patterns beyond the age effects (Sup Table 2). The most significant ones regarded the right hemisphere, particularly right AR with right IFG; right AF with right IFG/STG/AG; ACC with right AG. Besides, the asymmetry indices across pairs of tract and region were not related to each other (in the 3 groups of 6 comparisons r<0.40, p>0.42).

### Relationships between the structural and functional markers of maturation

To reduce the number of comparisons, we took into account the results of hemispheric asymmetries to further analyze the relationships between latencies, speeds and the microstructural properties of tracts and regions. When no asymmetry was observed, we considered averaged values (i.e. for contralateral P2 response, PT and HG). In the other cases (i.e. for measures showing asymmetries), we analyzed left and right values independently (i.e. for ipsilateral P2 responses, AR and AF tracts, IFG, MTG, STG and AG regions).

Similarly to our visual studies (Dubois et al. 2008, Adibpour, Dubois & Dehaene-Lambertz, 2018), we first evaluated the relationships between the microstructure of auditory tracts (λ_⊥_) and the conduction speed of P2 responses. Partial correlations controlling for age effects (Table 4a) were non-significant for the acoustic radiations and arcuate fascicles but showed a trend for the auditory callosal fibers and the left ipsilateral P2 response (Figure 5a). Although this result did not survive correction for multiple comparisons, it might suggest that the ipsilateral P2 response in the left hemisphere is modulated by the arrival of information from the right hemisphere through the callosal fibers, consistently with a previous independent study comparing infants with corpus callosum agenesis and typical infants (Adibpour, Dubois, Moutard, et al., 2018). Besides the asymmetry indices in P2 speeds and tracts microstructure were not related to each other (in the 2 groups of 3 comparisons r–0.42, p>0.32).

**Figure 5:**
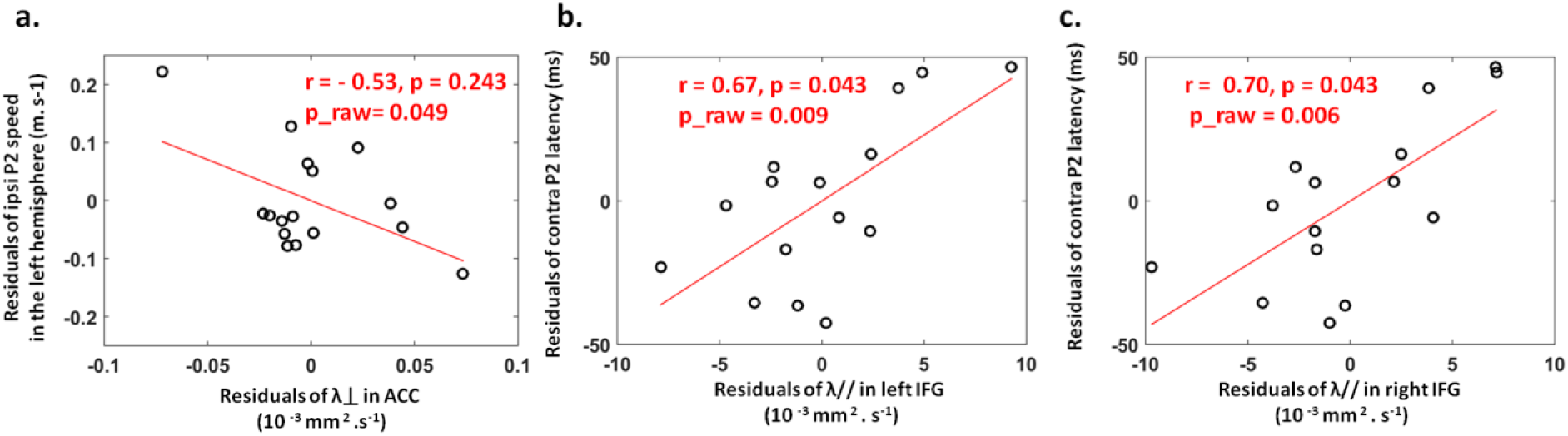
Relationships between P2 characteristics and the microstructural properties of the language network. a: The speed of ipsilateral P2 response in the left hemisphere tended to be related to transverse diffusivity (λ_⊥_) in the auditory callosal fibers after controlling for age effects (n=16, Table 3a). b/c: The latency of averaged contralateral responses was related to longitudinal diffusivity (λ_||_) in the left (b) and right IFG (c), after controlling for age effects (n=15, Table 3b). The plots represent the residuals in each parameter, after regressing out the effect of age. The partial correlation coefficients (r) and the corrected p values are indicated similarly to Table 3, as well as raw (uncorrected) p values.

**Table 3:**
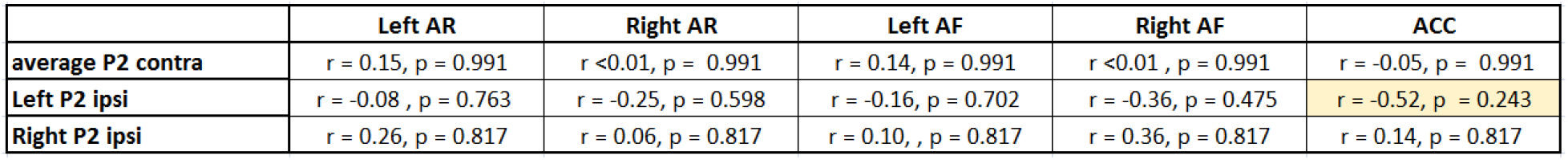
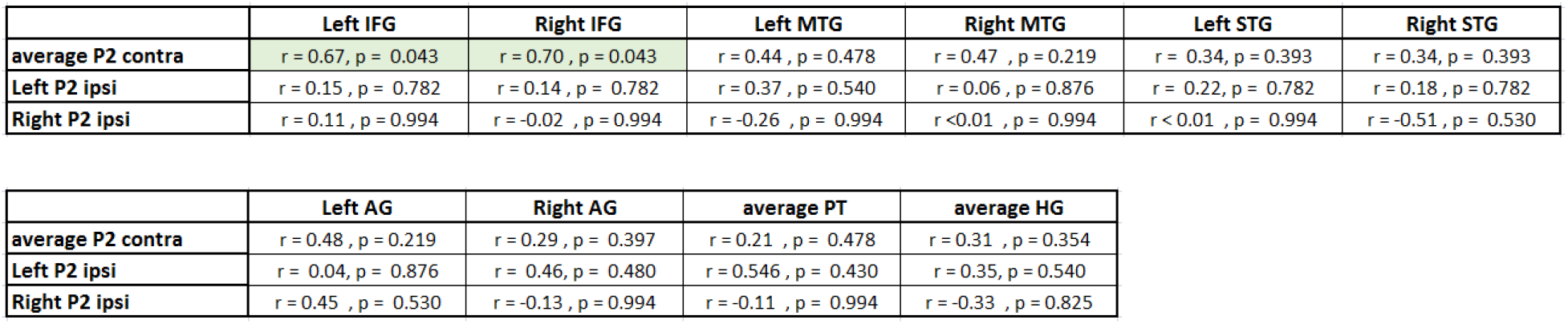
Relationships between the structural and functional markers of maturation. a: Correlation analyses between P2 speeds (contralateral responses averaged over both hemispheres; left and right ipsilateral responses) and the microstructure of auditory tracts (transverse diffusivity λ_⊥_ for left and right AR, AF and ACC), based on partial correlations taking into account the age effects (n=16). P-values were corrected for the number of comparisons made for each P2 response (c = 5) using FDR approach. The only observed effect was a trend for the speed of left ipsilateral response that correlated with the auditory callosal fibers (uncorrected p = 0.049, corrected p = 0.243 shaded in yellow, Figure 5a) but it did not survive the correction for multiple comparisons. b: Correlation analyses between P2 latencies (contralateral responses averaged over both hemispheres, as well as left and right ipsilateral responses) and the microstructure of perisylvian regions (longitudinal diffusivity λ_||_ for left and right regions when an asymmetry was observed, or averaged over both hemispheres), based on partial correlations taking into account the age effects (n=15). P-values were corrected for the number of comparisons made for each P2 latency (c=10) using FDR approach. The only significant relationships were between the latency of averaged contralateral response and the left and right inferior frontal gyri (p<0.05, shaded in green, Figure 5b,c). See Sup Tab 3 for similar analyses with P2 speeds.

|  | Left AR | Right AR | Left AF | Right AF | ACC |
| --- | --- | --- | --- | --- | --- |
| average P2 contra | r = 0.15, p = 0.991 | r <0.01, p = 0.991 | r = 0.14, p = 0.991 | r <0.01, p = 0.991 | r = -0.05, p = 0.991 |
| Left P2 ipsi | r = -0.08, p = 0.763 | r = -0.25, p = 0.598 | r = -0.16, p = 0.702 | r = -0.36, p = 0.475 | r = -0.52, p = 0.243 |
| Right P2 ipsi | r = 0.26, p = 0.817 | r = 0.06, p = 0.817 | r = 0.10, p = 0.817 | r = 0.36, p = 0.817 | r = 0.14, p = 0.817 |

|  | Left IFG | Right IFG | Left MTG | Right MTG | Left STG | Right STG |
| --- | --- | --- | --- | --- | --- | --- |
| average P2 contra | r = 0.67, p = 0.043 | r = 0.70, p = 0.043 | r = 0.44, p = 0.478 | r = 0.47, p = 0.219 | r = 0.34, p = 0.393 | r = 0.34, p = 0.393 |
| Left P2 ipsi | r = 0.15, p = 0.782 | r = 0.14, p = 0.782 | r = 0.37, p = 0.540 | r = 0.06, p = 0.876 | r = 0.22, p = 0.782 | r = 0.18, p = 0.782 |
| Right P2 ipsi | r = 0.11, p = 0.994 | r = -0.02, p = 0.994 | r = -0.26, p = 0.994 | r <0.01, p = 0.994 | r <0.01, p = 0.994 | r = -0.51, p = 0.530 |

|  | Left AG | Right AG | average PT | average HG |
| --- | --- | --- | --- | --- |
| average P2 contra | r = 0.48, p = 0.219 | r = 0.29, p = 0.397 | r = 0.21, p = 0.478 | r = 0.31, p = 0.354 |
| Left P2 ipsi | r = 0.04, p = 0.876 | r = 0.46, p = 0.480 | r = 0.546, p = 0.430 | r = 0.35, p = 0.540 |
| Right P2 ipsi | r = 0.45, p = 0.530 | r = -0.13, p = 0.994 | r = -0.11, p = 0.994 | r = -0.33, p = 0.825 |

Regarding the relationships between the microstructure of cortical regions (λ_//_) and P2 characteristics, we only observed links for the averaged latency of contralateral P2 responses with the left and right IFG (Table 4b, Figure 5b, c). When P2 speed was considered rather than latency, results were similar but less significant (Sup Table 3). Finally, the asymmetry indices in P2 latencies and regions microstructure were not related to each other (in the 2 groups of 6 comparisons: r<0.50, p>0.35).

## Discussion

In this study, we explored the maturation of the auditory system during the first postnatal semester both from a functional and a structural perspective. We showed that the auditory P2 response tends to be of higher amplitude in the left than right hemisphere following binaural stimulations, and in the contra-than ipsilateral hemisphere following monaural ones. We also observed longer latency for ipsi-than contralateral responses particularly in the left hemisphere, which might reflect an asymmetric inter-hemispheric transfer of responses. We further highlighted a number of microstructural asymmetries in acoustic radiations and arcuate fascicles for white matter, and in inferior frontal, angular, medial and superior temporal gyri for the cortex. Besides, we demonstrated intense developmental changes in P2 latencies, auditory tracts and perisylvian regions during this period. When evaluating the links between the functional and structural indices of maturation, we showed limited relationships between the response speeds and the microstructure of underlying white matter pathways: some trend was only observed between callosal fibers and left ipsilateral responses, consistently with a previous study (Adibpour et al, 2018). Finally, the latency of contralateral P2 responses was related to the microstructural properties of the inferior frontal gyri.

### Early markers of functional lateralization

Firstly, we studied the lateralization of P2 responses in two different auditory stimulation tasks in two groups of infants. Regarding P2 amplitude, we observed a weak lateralization in favor of the left hemisphere following the binaural presentation of syllables, suggesting a left dominance in processing consonant-vowel syllables. This is relatively coherent with previous functional MRI (fMRI) studies of young infants, in which activations to speech stimuli were reported to be larger in the left than right *planum temporale* or in the left temporal lobe in general, notably relative to non-linguistic stimuli (Dehaene-Lambertz, Dehaene, & Hertz-Pannier, 2002; Dehaene-Lambertz et al., 2010; Shultz et al., 2014).

In the monaural paradigm, we did not find any left-right difference for P2 amplitude, perhaps because this was alleviated by the strong contra-laterality effect that we observed whatever the ear of stimulation. We also suspect that P2 responses and their lateralization might have been modulated by the orientation of attention to the sides of the auditory space. Indeed, the side of syllable presentation remained constant within each block of trials, allowing infants to direct their attention towards one ear or the other. In dichotic listening tasks where subjects are asked to report the perceived syllable, reports from the right ear are generally more accurate than the left ear (this is known as the right ear advantage). However, in these tasks, overt attention to a given side alters perceptual performances such that the right ear advantage is increased when the right ear is attended, and decreased or even reversed when the left ear is attended (Hugdahl & Westerhausen, 2016). Therefore, responses in the context of the binaural paradigm (when spatial attention was not solicited) might better represent the intrinsic lateralization in our infant studies.

Regarding P2 latency, we observed a delay in ipsilateral compared to contralateral responses when syllables were presented monaurally, whereas in a previous study (Adibpour, Dubois, Moutard, et al., 2018), such a delay was observed only when the stimulus (“babble noise” corresponding to overlaid segments of speech) was presented in the left ear. This difference might be explained by experimental variations: in the latter study, the side of presentation was randomly chosen at each trial while here, the side of the presentation alternated across blocks of 48 trials, probably leading to sensory priming within blocks.

Finally, we observed a delay in the left ipsilateral responses compared to the right equivalent. It is consistent with our previous study showing a similar asymmetric pattern of responses in another group of typical infants of the same age (Adibpour, Dubois, Moutard, et al., 2018). However, because this delay was not seen in infants with corpus callosum agenesis, we attributed it to an inter-hemispheric transfer of information via the callosal connections. The spatial resolution of EEG cannot disentangle the direct ipsilateral and transcallosal responses, which progressively merge and thus might delay the P2 peak that we measured. Because this delay is seen only for the left ipsilateral response and not the right, it suggests an asymmetric transfer of information, prominent from the right to the left hemisphere. A recent study in adults also described asymmetry in the auditory inter-hemispheric connectivity: TMS applied over the right (but not the left) auditory regions changes the functional connectivity between these regions during resting-state, in proportion with the volume of auditory callosal fibers (Andoh, Matsushita, & Zatorre, 2015). Similarly, some studies have reported broader inter-hemispheric functional connectivity for the regions of the right hemisphere, but stronger intra-hemispheric connectivity for the left hemisphere, especially for linguistic areas (Gotts et al., 2013). Our two studies in infants thus suggest that the asymmetric pattern of inter-hemispheric connectivity is already observed in early development.

### Early structural asymmetries between hemispheres

Acoustic radiations and arcuate fascicles showed lower diffusivities in the left than right tracts, suggesting more complex, denser or more mature patterns of microstructure in the left than right tracts. Acoustic radiations have not been studied in infants so far, but leftward DTI asymmetries have already been reported for the arcuate fasciculus during the first postnatal semester, both in its structure and its faster left maturation (Dubois et al., 2009; Dubois, Poupon, et al., 2016). In adults, previous *post-mortem* findings have reported more myelinated axons in the left than in the right posterior superior temporal lobe, an area posterior to acoustic radiations (Anderson, Southern, & Powers, 1999). Nevertheless, our results should be interpreted with caution because the acoustic radiations are quite short in length, and they partially cross the optic radiations in their path between the thalamus and the auditory cortex. Although their reconstruction with probabilistic tractography (Behrens et al., 2007) was feasible despite infants’ immature fibers, it was still challenging because of the limited spatial resolution relatively to infant brain size. Thus, our DTI parameters might be prone to partial volume effects with other tracts. Further work is needed to confirm the presence of this acoustic radiation asymmetry in infants and investigate how it relates to other regional asymmetries.

Asymmetries in the perisylvian cortical areas were observed in favor of both hemispheres, with more advanced maturation (lower longitudinal diffusivity) relatively to their homologues, in the right hemisphere for inferior frontal and superior temporal gyri, and in the left hemisphere for angular and middle temporal gyri. These observations are partly in agreement with previous findings based on T2w intensity, in which we showed rightward STS asymmetry (Leroy et al, 2011). This latter study was limited to the sulci whereas here, parameters were averaged across relatively large parcels covering the gyri. These DTI asymmetries might be explained by two possible factors. On one hand, homologous tissues might have differences in cells and organelles density from birth on, due to genetic or epigenetic factors, and this would extend throughout lifespan. For example, the columns are wider and with more neuropil space in the left than right *planum temporale* in the adult brain (Buxhoeveden, Switala, Litaker, Roy, & Casanova, 2001), which certainly impacts DTI parameters. On the other hand, these differences might be due to transient asynchronies in left-right maturation engendering a different development of dendritic arborization and intra-cortical myelination (i.e. faster maturation in a region of one hemisphere relatively to its counterpart). Distinguishing these two aspects would require studying infants relatively to an adult population, as we have recently done to compare the maturation of dorsal and ventral linguistic pathways throughout infancy (Dubois et al., 2016). Such an analysis was not possible here. In any case, the difference in hemispheric asymmetries between the superior and middle temporal gyri would need to be further explored as it could be one of the relevant markers distinguishing general auditory vs. linguistic regions.

### Maturational evolution of functional and structural auditory markers

As expected from the literature (Barnet et al., 1975; Novak et al., 1989; Kushnerenko et al., 2002; for a review see Wunderlich & Cone-Wesson., 2006), P2 latency decreased with the infants’ increasing age in all experimental conditions. This suggests an improvement in functional efficiency over the first postnatal months. As detailed in the last section of the discussion, numerous processes along the auditory pathways probably contribute to this improvement (e.g. maturation of cochlea, brainstem nuclei, auditory cortices and other cortical sources, as well as the myelination of auditory white matter pathways).

In the white matter (acoustic radiations, arcuate fascicles and callosal fibers), transverse diffusivity consistently decreased with the infants’ age, suggesting intense myelination of the pathways (Dubois, Dehaene-Lambertz, et al., 2014; Yakovlev. & Lecours., 1967). In the grey matter, longitudinal diffusivity also decreased with infants’ age. Various developmental mechanisms involved in grey matter maturation at this age might underly these changes, including the development of dendritic arborization (Ball et al., 2013), the increase in synaptic density, the myelination of intra-cortical fibers (Lebenberg, Labit, et al., 2019) or the changes in intracellular organelle density (for a review see (Ouyang, Dubois, Yu, Mukherjee, & Huang, 2019)). A recent study using another diffusion-based method (neurite orientation dispersion and density imaging: NODDI) also pointed to the increasing cellular and organelle densities around the term period as sources of developmental changes in diffusion parameters (Batalle et al., 2019). Overall, our results confirmed that tracts and regions were at different microstructural stages, which was consistent with the *post-mortem* and *in vivo* descriptions of maturational progression: associative white matter pathways lag behind the sensory projection pathways (Yakovlev. & Lecours., 1967), and multi-modal cortices lag behind the primary ones (Brody, Kinney, Kloman, & Gilles, 1987; Leroy et al., 2011). Nevertheless, as discussed above, these tracts and regions have also different microstructural organization at the mature stage (Dubois, Dehaene-Lambertz, et al., 2014; Dubois, Poupon, et al., 2016; Fukutomi et al., 2018). Thus, interpreting DTI differences in terms of maturation should be done cautiously without pondering over adult measures.

### Relationships between the functional and structural markers of maturation

MRI data were acquired only for the second EEG group of infants, limiting the number of structure-function relationships that we could analyze. Although we might have expected that the functional lateralization in auditory responses was dependent on structural asymmetries, we did not observe such a relationship. In particular, the asymmetry in AR microstructure did not translate in P2 latency differences across hemispheres, perhaps because this late component is less sensitive to AR inputs than earlier components. However, such early peaks are difficult to identify in each infant with EEG. In the future, MEG might be a more sensitive technique to be considered for such purposes. Regarding asymmetries in cortical microstructure, the leftward asymmetry of the angular gyrus might explain the larger fMRI activations reported in 3-months old infants listening to speech (Dehaene-Lambertz et al., 2002).

Regarding developmental changes, we first expected that the myelination of auditory pathways would increase P2 speed, in particular for the acoustic radiations since their myelination starts after birth and progresses slowly until 3-4 years of age (Yakovlev & Lecours, 1974). Nevertheless, we did not observe any direct relationship, contrarily to our findings in the visual system that highlighted some for P1 speed and optic radiations, P1 interhemispheric transfer and visual callosal fibers (Adibpour, Dubois, & Dehaene-Lambertz, 2018; Dubois et al., 2008). Actually, auditory P2 is a relatively late cortical response that probably arises from computations engaging multiple generators varying with the task and maturing in a complex way. In support of this view, EEG modelling approaches of P2 responses evoked by vowels and tones have proposed cortical sources in the superior temporal, supra-marginal and inferior frontal gyri in two-month-old infants (Basirat, Dehaene, & Dehaene-Lambertz, 2014; Bristow et al., 2009), or in the anterior cingulate and temporal regions for 6-month-olds (Ortiz-Mantilla, Hämäläinen, & Benasich, 2012). Imada et al. (2006) also reported superior temporal and inferior frontal sources using MEG in 6 and 12 month-olds (Imada et al., 2006). These studies hypothesized that inferior frontal regions are involved in syllable perception, which might explain the relationship we observed between the latency of P2 contralateral responses and the IFG microstructure. On the other hand, a recent study of the aging population suggested that structure-function relationships differ between the visual and auditory modalities (Price et al., 2017). For this latter one, cumulative delays in evoked responses were related to differences in the gray matter volume of superior temporal regions, but not in auditory tracts (Price et al., 2017). This highlights the importance of considering cortical properties when analyzing the inter-individual variability in auditory functional responses.

In the light of our previous study comparing typical infants and infants with agenesis of corpus callosum (Adibpour, Dubois, Moutard, et al., 2018), we also assumed a possible role of auditory callosal fibers in the ERP characteristics, particularly for ipsilateral responses measured in the left hemisphere. Although it should be confirmed over a larger cohort of infants, the trend we observed between the callosal microstructure and the speed of this response is coherent with our hypothesis. Previous studies in adults further suggested that while the right-ear stimuli can access the left hemisphere processing resources through the strong contralateral auditory pathway, left-ear stimuli might access these resources through callosal connections from the right to the left hemisphere. Westerhausen et al. (2008) demonstrated that the accuracy of reports from the left, but not the right ear, is related to the strength of inter-hemispheric connections between superior temporal regions (Westerhausen, Grüner, Specht, & Hugdahl, 2008). Left-ear reports in dichotic paradigm are also disrupted in patients with commissurotomy (Milner, Taylor, & Sperry, 1968; Sparks & Geschwind, 1968) and in patients with lesions to posterior part of corpus callosum (Pollmann, Maertens, von Cramon, Lepsien, & Hugdahl, 2002).

All together, these results show that combining dedicated EEG and MRI approaches in infants can shed light on the complex maturational relationships between the functional responses to auditory stimuli and the properties of the underlying neural network that involves both white matter pathways and cortical gray matter regions.

## Conclusion

This study highlighted the early lateralization and the developmental processes occurring within the auditory system during the first semester of infancy. Compared with the visual modality, our results taken together suggested complex functional and structural patterns and relationships. Auditory P2 responses might depend both on callosal connectivity and on the maturation of inferior frontal gyri. Yet, considering earlier responses (measured with MEG for example) would help us to understand the role of auditory projection pathways and primary regions. Measuring other complementary MRI measures (e.g. T1 relaxation time (Lebenberg et al, 2019)) would also provide a more comprehensive evaluation of the auditory network, and of the ongoing maturational processes. Lastly, the detailed comparison of different brain systems showing asynchronous maturation would offer critical cues for the understanding of human cognitive development.

## Acknowledgements

This research was supported by grants from the Fondation de France and the Fyssen Foundation. The funders had no role in study design, data collection and analysis, decision to publish, or preparation of the manuscript. The authors would like to thank all the infants and their parents who participated in this study as well as Giovanna Santoro and the medical team of UNIACT at Neurospin, who helped in carrying out the experiments. We are also grateful to Francois Leroy for his help in MRI analyses.

The authors have no conflict of interest to declare.

---

**Supplementary Table 1.** Relationships between the microstructure of auditory tracts (a) or perisylvian regions (b). Partial correlations were computed between transverse diffusivity (λ_⊥_) values in pairs of white matter tracts (a, n=22), or between longitudinal diffusivity (λ_||_) values in pairs of cortical regions (b, n=21) while controlling for age to remove common dependencies. P-values were corrected for the number of comparisons made within each hemisphere (c=3 for tracts, c=15 for regions) and across hemispheres (c=2 for tracts, c=6 for regions) using FDR approach. Significant (p<0.05) and marginally significant (p<0.1) comparisons are shaded in green and yellow respectively.

| Left hemisphere | AF | ACC |
| --- | --- | --- |
| AR | $r = 0.66, p = 0.011$ | $r = 0.33, p = 0.272$ |
| AF | | $r = 0.54, p = 0.046$ |
| Right hemisphere | AF | ACC |
| AR | $r = 0.21, p = 0.473$ | $r = 0.19, p = 0.473$ |
| AF | | $r = 0.69, p = 0.007$ |
| Across Hemispheres | Right AR | Right AF |
| Left AR | $r = 0.63, p = 0.007$ | |
| Left AF | | $r = 0.76, p < 0.001$ |

b:
| Left Hemisphere | IFG | MTG | STG | HG | PT | AG |
| --- | --- | --- | --- | --- | --- | --- |
| IFG | | $r = 0.46, p = 0.064$ | $r = 0.80, p < 0.001$ | $r = 0.17, p = 0.550$ | $r = 0.46, p = 0.064$ | $r = 0.51, p = 0.049$ |
| MTG | | | $r = 0.78, p < 0.001$ | $r = 0.35, p = 0.161$ | $r = 0.49, p = 0.049$ | $r = 0.50, p = 0.049$ |
| STG | | | | $r = 0.42, p = 0.085$ | $r = 0.56, p = 0.042$ | $r = 0.55, p = 0.042$ |
| HG | | | | | $r = 0.50, p = 0.049$ | $r = 0.08, p = 0.742$ |
| PT | | | | | | $r = 0.12, p = 0.646$ |
| AG |  |  |  |  |  |  |
| Right Hemisphere | IFG | MTG | STG | HG | PT | AG |
| IFG | | $r = 0.58, p = 0.017$ | $r = 0.73, p < 0.001$ | $r = 0.62, p = 0.016$ | $r = 0.27, p = 0.256$ | $r = 0.60, p = 0.016$ |
| MTG | | | $r = 0.47, p = 0.060$ | $r = 0.57, p = 0.019$ | $r = 0.33, p = 0.184$ | $r = 0.63, p = 0.016$ |
| STG | | | | $r = 0.60, p = 0.016$ | $r = 0.27, p = 0.256$ | $r = 0.52, p = 0.037$ |
| HG | | | | | $r = 0.36, p = 0.167$ | $r = 0.33, p = 0.184$ |
| PT | | | | | | $r = 0.36, p = 0.167$ |
| AG |  |  |  |  |  |  |
| Across Hemispheres | Right IFG | Right MTG | Right STG | Right HG | Right PT | Right AG |
| Left IFG | $r = 0.82, p < 0.001$ | | | | | |
| Left MTG | | $r = 0.46, p = 0.067$ | | | | |
| Left STG | | | $r = 0.55, p = 0.034$ | | | |
| Left HG | | | | $r = 0.15, p = 0.583$ | | |
| Left PT | | | | | $r = 0.13, p = 0.583$ | |
| Left AG | | | | | | $r = 0.45, p = 0.067$ |

**Supplementary Table 2.** Relationships between the microstructure of auditory tracts and perisylvian regions. Partial correlations were computed between pairs of tract and region within each hemisphere (considering λ_⊥_ values for tracts, λ_||_ values for regions), while controlling for age effects to remove common dependences (n=21). P-values were corrected for the number of comparisons made within each hemisphere (c=18) using FDR approach. Significant (p<0.05) and marginally significant (p<0.1) comparisons are shaded in green and yellow.

| Left Hemisphere | IFG | MTG | STG | HG | PT | AG |
| --- | --- | --- | --- | --- | --- | --- |
| AR | r = 0.37, p = 0.253 | r = 0.31, p = 0.356 | r = 0.44, p = 0.253 | r = 0.13, p = 0.608 | r = 0.37, p = 0.253 | r = -0.05, p = 0.827 |
| AF | r = 0.37, p = 0.253 | r = 0.57, p = 0.074 | r = 0.58, p = 0.074 | r = 0.29, p = 0.395 | r = 0.37, p = 0.253 | r = 0.14, p = 0.608 |
| ACC | r = 0.14, p = 0.608 | r = 0.41, p = 0.253 | r = 0.27, p = 0.410 | r = 0.24, p = 0.436 | r = 0.24, p = 0.436 | r = 0.14, p = 0.608 |
| Right hemisphere | IFG | MTG | STG | HG | PT | AG |
| AR | r = 0.58, p = 0.044 | r = 0.47, p = 0.079 | r = 0.44, p = 0.089 | r = 0.36, p = 0.165 | r = 0.10, p = 0.684 | r = 0.46, p = 0.079 |
| AF | r = 0.66, p = 0.015 | r = 0.46, p = 0.079 | r = 0.56, p = 0.048 | r = 0.51, p = 0.067 | r = 0.26, p = 0.284 | r = 0.66, p = 0.015 |
| ACC | r = 0.27, p = 0.284 | r = 0.36, p = 0.165 | r = 0.28, p = 0.284 | r = 0.26, p = 0.283 | r = 0.45, p = 0.087 | r = 0.54, p = 0.051 |

**Supplementary Table 3.** Relationships between the microstructure of auditory tracts and perisylvian regions and the speed of P2 responses. Correlations between P2 speeds (contralateral responses averaged over both hemispheres, as well as left and right ipsilateral responses) and the microstructure of perisylvian regions (longitudinal diffusivity λ_||_ for left and right regions when an asymmetry was observed, or averaged over both hemispheres), based on partial correlations taking into account the age effects (n=15). As no correlation survived the correction for multiple comparisons (c=10 for each P2 speed), we here show uncorrected p-values (p<0.05 and p<0.1 are shaded in green and yellow respectively). Note that the general pattern of correlations from table 3. b was replicated, with comparable correlation coefficients but less significance.

|  | Left IFG | Right IFG | Left MTG | Right MTG | Left STG | Right STG |
| --- | --- | --- | --- | --- | --- | --- |
| average P2 contra | r = -0.46, p = 0.096 | r = -0.51, p = 0.059 | r = -0.16, p = 0.594 | r = -0.43, p = 0.121 | r = -0.14, p = 0.639 | r = -0.25, p = 0.386 |
| Left P2 ipsi | r = -0.07, p = 0.813 | r = -0.11, p = 0.705 | r = -0.45, p = 0.105 | r = -0.14, p = 0.641 | r = -0.22, p = 0.441 | r = -0.33, p = 0.249 |
| Right P2 ipsi | r = 0.01, p = 0.975 | r = 0.10, p = 0.719 | r = -0.08, p = 0.777 | r = 0.33, p = 0.241 | r = 0.10, p = 0.737 | r = 0.50, p = 0.070 |

|  | Left AG | Right AG | average PT | average HG |
| --- | --- | --- | --- | --- |
| average P2 contra | r = -0.40, p = 0.153 | r = -0.20, p = 0.483 | r = -0.19, p = 0.520 | r = -0.26, p = 0.362 |
| Left P2 ipsi | r = -0.002, p = 0.993 | r = -0.42, p = 0.129 | r = -0.58, p = 0.029 | r = -0.45, p = 0.103 |
| Right P2 ipsi | r = -0.43, p = 0.122 | r = 0.14, p = 0.633 | r = 0.06, p = 0.847 | r = 0.28, p = 0.325 |

## References

Adibpour, P., Dubois, J., & Dehaene-Lambertz, G. (2018). Right but not left hemispheric discrimination of faces in infancy. Nature Human Behaviour, 2(1), 67.

Adibpour, P., Dubois, J., Moutard, M.-L., & Dehaene-Lambertz, G. (2018). Early asymmetric inter-hemispheric transfer in the auditory network: insights from infants with corpus callosum agenesis. Brain Structure and Function, 1–13.

Anderson, B., Southern, B. D., & Powers, R. E. (1999). Anatomic asymmetries of the posterior superior temporal lobes: a postmortem study. Cognitive and Behavioral Neurology, 12(4), 247–254.

Andoh, J., Matsushita, R., & Zatorre, R. J. (2015). Asymmetric interhemispheric transfer in the auditory network: evidence from TMS, resting-state fMRI, and diffusion imaging. Journal of neuroscience, 35(43), 14602–14611.

Ball, G., Srinivasan, L., Aljabar, P., Counsell, S. J., Durighel, G., Hajnal, J. V., … Edwards, A.D. (2013). Development of cortical microstructure in the preterm human brain. Proc Natl Acad Sci U S A, 110(23), 9541–9546. doi:10.1073/pnas.1301652110

Barnet, A. B., Ohlrich, E. S., Weiss, I. P., & Shanks, B. (1975). Auditory evoked potentials during sleep in normal children from ten days to three years of age. Electroencephalography and clinical neurophysiology, 39(1), 29–41.

Basirat, A., Dehaene, S., & Dehaene-Lambertz, G. (2014). A hierarchy of cortical responses to sequence violations in three-month-old infants. Cognition, 132(2), 137–150. doi:10.1016/j.cognition.2014.03.013

Batalle, D., O’Muircheartaigh, J., Makropoulos, A., Kelly, C. J., Dimitrova, R., Hughes, E. J., … Edwards, A.D. (2019). Different patterns of cortical maturation before and after 38 weeks gestational age demonstrated by diffusion MRI in vivo. Neuroimage.

Boemio, A., Fromm, S., Braun, A., & Poeppel, D. (2005). Hierarchical and asymmetric temporal sensitivity in human auditory cortices. Nat Neurosci, 8(3), 389–395.

Bristow, D., Dehaene-Lambertz, G., Mattout, J., Soares, C., Gliga, T., Baillet, S., & Mangin, J. F. (2009). Hearing faces: how the infant brain matches the face it sees with the speech it hears. J Cogn Neurosci, 21(5), 905–921.

Brody, B. A., Kinney, H. C., Kloman, A. S., & Gilles, F. H. (1987). Sequence of central nervous system myelination in human infancy. I. An autopsy study of myelination. J Neuropathol Exp Neurol, 46(3), 283–301.

Buxhoeveden, D. P., Switala, A. E., Litaker, M., Roy, E., & Casanova, M. F. (2001). Lateralization of minicolumns in human planum temporale is absent in nonhuman primate cortex. Brain, Behavior and Evolution, 57(6), 349–358.

Dehaene-Lambertz, G., Dehaene, S., & Hertz-Pannier, L. (2002). Functional neuroimaging of speech perception in infants. Science, 298(5600), 2013–2015. doi:10.1126/science.1077066

Dehaene-Lambertz, G., Montavont, A., Jobert, A., Allirol, L., Dubois, J., Hertz-Pannier, L., & Dehaene, S. (2010). Language or music, mother or Mozart? Structural and environmental influences on infants’ language networks. Brain and language, 114(2), 53–65.

Dubois, J., Adibpour, P., Poupon, C., Hertz-Pannier, L., & Dehaene-Lambertz, G. (2016). MRI and M/EEG studies of the white matter development in human fetuses and infants: review and opinion. Brain Plasticity, 2(1), 49–69.

Dubois, J., Benders, M., Lazeyras, F., Borradori-Tolsa, C., Leuchter, R. H., Mangin, J. F., & Huppi, P. S. (2010). Structural asymmetries of perisylvian regions in the preterm newborn. Neuroimage, 52(1), 32–42. doi:10.1016/j.neuroimage.2010.03.054

Dubois, J., Dehaene-Lambertz, G., Kulikova, S., Poupon, C., Huppi, P. S., & Hertz-Pannier, L. (2014). The early development of brain white matter: a review of imaging studies in fetuses, newborns and infants. Neuroscience, 276, 48–71. doi:10.1016/j.neuroscience.2013.12.044

Dubois, J., Dehaene-Lambertz, G., Soares, C., Cointepas, Y., Le Bihan, D., & Hertz-Pannier, L. (2008). Microstructural correlates of infant functional development: example of the visual pathways. J Neurosci, 28(8), 1943–1948. doi:10.1523/JNEUROSCI.5145-07.2008

Dubois, J., Hertz-Pannier, L., Cachia, A., Mangin, J. F., Le Bihan, D., & Dehaene-Lambertz, G. (2009). Structural asymmetries in the infant language and sensori-motor networks. Cereb Cortex, 19(2), 414–423. doi:10.1093/cercor/bhn097

Dubois, J., Hertz-Pannier, L., Dehaene-Lambertz, G., Cointepas, Y., & Le Bihan, D. (2006). Assessment of the early organization and maturation of infants’ cerebral white matter fiber bundles: a feasibility study using quantitative diffusion tensor imaging and tractography. Neuroimage, 30(4), 1121–1132.

Dubois, J., Kulikova, S., Hertz-Pannier, L., Mangin, J. F., Dehaene-Lambertz, G., & Poupon, C. (2014). Correction strategy for diffusion-weighted images corrupted with motion: application to the DTI evaluation of infants’ white matter. Magn Reson Imaging, 32(8), 981–992. doi:10.1016/j.mri.2014.05.007

Dubois, J., Lefèvre, J., Angleys, H., Leroy, F., Fischer, C., Lebenberg, J., … Hertz-Pannier, L. (2019). The dynamics of cortical folding waves and prematurity-related deviations revealed by spatial and spectral analysis of gyrification. Neuroimage.

Dubois, J., Poupon, C., Thirion, B., Simonnet, H., Kulikova, S., Leroy, F., … Dehaene-Lambertz, G. (2016). Exploring the Early Organization and Maturation of Linguistic Pathways in the Human Infant Brain. Cereb Cortex, 26(5), 2283–2298. doi:10.1093/cercor/bhv082

Duclap, D., Schmitt, B., Riff, O., Guevara, P., Marrakchi-Kacem, L., Brion, V., … Poupon, C. (2012). Connectomist-2.0: a novel diffusion analysis toolbox for BrainVISA. Paper presented at the Paper presented at the 29th ESMRMB, Lisbone, Portugal.

Flechsig, P. E. (1920). Anatomie des menschlichen Gehirns und Rückenmarks auf myelogenetischer Grundlage (Vol. 1): G. Thieme.

Fukutomi, H., Glasser, M. F., Zhang, H., Autio, J. A., Coalson, T. S., Okada, T., … Hayashi, T. (2018). Neurite imaging reveals microstructural variations in human cerebral cortical gray matter. Neuroimage.

Glasel, H., Leroy, F., Dubois, J., Hertz-Pannier, L., Mangin, J. F., & Dehaene-Lambertz, G. (2011). A robust cerebral asymmetry in the infant brain: the rightward superior temporal sulcus. Neuroimage, 58(3), 716–723. doi:10.1016/j.neuroimage.2011.06.016

Gotts, S. J., Jo, H. J., Wallace, G. L., Saad, Z. S., Cox, R. W., & Martin, A. (2013). Two distinct forms of functional lateralization in the human brain. Proc Natl Acad Sci U S A, 110(36), E3435–3444. doi:10.1073/pnas.1302581110

Hugdahl, K., & Westerhausen, R. (2016). Speech processing asymmetry revealed by dichotic listening and functional brain imaging. Neuropsychologia, 93, 466–481.

Huppi, P. S., Maier, S. E., Peled, S., Zientara, G. P., Barnes, P. D., Jolesz, F. A., & Volpe, J. J. (1998). Microstructural development of human newborn cerebral white matter assessed in vivo by diffusion tensor magnetic resonance imaging. Pediatr Res, 44(4), 584–590. doi:10.1203/00006450-199810000-00019

Imada, T., Zhang, Y., Cheour, M., Taulu, S., Ahonen, A., & Kuhl, P. K. (2006). Infant speech perception activates Broca’s area: a developmental magnetoencephalography study. Neuroreport, 17(10), 957–962.

Kabdebon, C., Leroy, F., Simmonet, H., Perrot, M., Dubois, J., & Dehaene-Lambertz, G. (2014). Anatomical correlations of the international 10-20 sensor placement system in infants. Neuroimage, 99, 342–356. doi:10.1016/j.neuroimage.2014.05.046

Kushnerenko, E., Ceponiene, R., Balan, P., Fellman, V., Huotilainen, M., & Näätänen, R. (2002). Maturation of the auditory event-related potentials during the first year of life. Neuroreport, 13(1), 47–51.

Lebenberg, J., Labit, M., Auzias, G., Mohlberg, H., Fischer, C., Rivière, D., … Labra, N. (2019). A framework based on sulcal constraints to align preterm, infant and adult human brain images acquired in vivo and post mortem. Brain Structure and Function, 1–16.

Lebenberg, J., Mangin, J. F., Thirion, B., Poupon, C., Hertz-Pannier, L., Leroy, F., … Dubois, J. (2019). Mapping the asynchrony of cortical maturation in the infant brain: A MRI multi-parametric clustering approach. Neuroimage. doi:10.1016/j.neuroimage.2018.07.022

Lebenberg, J., Poupon, C., Thirion, B., Leroy, F., Mangin, J.-F., Dehaene-Lambertz, G., & Dubois, J. (2015). Clustering the infant brain tissues based on microstructural properties and maturation assessment using multi-parametric MRI. Paper presented at the Biomedical Imaging (ISBI), 2015 IEEE 12th International Symposium on.

Leroy, F., Glasel, H., Dubois, J., Hertz-Pannier, L., Thirion, B., Mangin, J. F., & Dehaene-Lambertz, G. (2011). Early maturation of the linguistic dorsal pathway in human infants. J Neurosci, 31(4), 1500–1506. doi:10.1523/JNEUROSCI.4141-10.2011

Li, G., Lin, W., Gilmore, J. H., & Shen, D. (2015). Spatial patterns, longitudinal development, and hemispheric asymmetries of cortical thickness in infants from birth to 2 years of age. Journal of neuroscience, 35(24), 9150–9162.

Lippé, S., Kovacevic, N., & McIntosh, A. R. (2009). Differential maturation of brain signal complexity in the human auditory and visual system. Frontiers in Human Neuroscience, 3.

Mahmoudzadeh, M., Dehaene-Lambertz, G., Fournier, M., Kongolo, G., Goudjil, S., Dubois, J., … Wallois, F. (2013). Syllabic discrimination in premature human infants prior to complete formation of cortical layers. Proc Natl Acad Sci U S A, 110(12), 4846–4851. doi:1212220110[pii]10.1073/pnas.1212220110

McCulloch, D. L., Orbach, H., & Skarf, B. (1999). Maturation of the pattern-reversal VEP in human infants: a theoretical framework. Vision Res, 39(22), 3673–3680.

Milner, B., Taylor, L., & Sperry, R. W. (1968). Lateralized suppression of dichotically presented digits after commissural section in man. Science, 161(3837), 184–185.

Monson, B. B., Eaton-Rosen, Z., Kapur, K., Liebenthal, E., Brownell, A., Smyser, C. D., … Neil, J.J. (2018). Differential Rates of Perinatal Maturation of Human Primary and Nonprimary Auditory Cortex. eNeuro, 5(1). doi:10.1523/ENEURO.0380-17.2017

Novak, G. P., Kurtzberg, D., Kreuzer, J. A., & Vaughan, H. G. (1989). Cortical responses to speech sounds and their formants in normal infants: maturational sequence and spatiotemporal analysis. Electroencephalography and clinical neurophysiology, 73(4), 295–305.

Ortiz-Mantilla, S., Hämäläinen, J. A., & Benasich, A. A. (2012). Time course of ERP generators to syllables in infants: a source localization study using age-appropriate brain templates. Neuroimage, 59(4), 3275–3287.

Ouyang, M., Dubois, J., Yu, Q., Mukherjee, P., & Huang, H. (2019). Delineation of early brain development from fetuses to infants with diffusion MRI and beyond. Neuroimage.

Perani, D., Saccuman, M. C., Scifo, P., Anwander, A., Spada, D., Baldoli, C., … Friederici, A.D. (2011). Neural language networks at birth. Proceedings of the National Academy of Sciences, 108(38), 16056–16061.

Pollmann, S., Maertens, M., von Cramon, D. Y., Lepsien, J., & Hugdahl, K. (2002). Dichotic listening in patients with splenial and nonsplenial callosal lesions. Neuropsychology, 16(1), 56.

Ponton, C. W., Eggermont, J. J., Kwong, B., & Don, M. (2000). Maturation of human central auditory system activity: evidence from multi-channel evoked potentials. Clinical Neurophysiology, 111(2), 220–236.

Price, D., Tyler, L. K., Henriques, R. N., Campbell, K., Williams, N., Treder, M., … Calder, A.C. (2017). Age-related delay in visual and auditory evoked responses is mediated by white-and grey-matter differences. Nature communications, 8, 15671.

Roberts, T. P., Khan, S. Y., Blaskey, L., Dell, J., Levy, S. E., Zarnow, D. M., & Edgar, J. C. (2009). Developmental correlation of diffusion anisotropy with auditory-evoked response. Neuroreport, 20(18), 1586–1591.

Rolland, C., Lebenberg, J., Leroy, F., Moulton, E., Adibpour, P., Rivière, D., … Dubois, J. (2019). Exploring microstructure asymmetries in the infant brain cortex: A methodological framework combining structural and diffusion MRI. Paper presented at the IEEE ISBI, Venice, Italy.

Shafer, V. L., Yan, H. Y., & Wagner, M. (2015). Maturation of cortical auditory evoked potentials (CAEPs) to speech recorded from frontocentral and temporal sites: three months to eight years of age. International Journal of Psychophysiology, 95(2), 77–93.

Shultz, S., Vouloumanos, A., Bennett, R. H., & Pelphrey, K. (2014). Neural specialization for speech in the first months of life. Dev Sci, 17(5), 766–774. doi:10.1111/desc.12151

Song, S. K., Sun, S. W., Ju, W. K., Lin, S. J., Cross, A. H., & Neufeld, A. H. (2003). Diffusion tensor imaging detects and differentiates axon and myelin degeneration in mouse optic nerve after retinal ischemia. Neuroimage, 20(3), 1714–1722.

Song, S. K., Yoshino, J., Le, T. Q., Lin, S. J., Sun, S. W., Cross, A. H., & Armstrong, R. C. (2005). Demyelination increases radial diffusivity in corpus callosum of mouse brain. Neuroimage, 26(1), 132–140. doi:10.1016/j.neuroimage.2005.01.028

Sowell, E. R., Peterson, B. S., Thompson, P. M., Welcome, S. E., Henkenius, A. L., & Toga, A. W. (2003). Mapping cortical change across the human life span. Nat Neurosci, 6(3), 309–315. doi:10.1038/nn1008

Sparks, R., & Geschwind, N. (1968). Dichotic listening in man after section of neocortical commissures. Cortex, 4(1), 3–16.

Thomas, J. L., Spassky, N., Perez Villegas, E. M., Olivier, C., Cobos, I., Goujet-Zalc, C., … Zalc, B. (2000). Spatiotemporal development of oligodendrocytes in the embryonic brain. J Neurosci Res, 59(4), 471–476. doi:10.1002/(SICI)1097-4547(20000215)59:4<471::AID-JNR1>3.0.CO;2-3

Westerhausen, R., Grüner, R., Specht, K., & Hugdahl, K. (2008). Functional relevance of interindividual differences in temporal lobe callosal pathways: a DTI tractography study. Cerebral cortex, 19(6), 1322–1329.

Wunderlich, J. L., & Cone-Wesson, B. K. (2006). Maturation of CAEP in infants and children: a review. Hear Res, 212(1-2), 212–223. doi:10.1016/j.heares.2005.11.008

Yakovlev, P. L., & Lecours, A. R. (1967). (Vol. in Regional Development of the Brain in Early Life (ed. Minkowski, A.)): Blackwell, Oxford.

Yakovlev., & Lecours. (1967). The myelogenetic cycles of regional maturation in the brain.: Oxford: Blackwell.

